# ADToolbox: Incorporating Metagenomics Data for Improved Prediction of Anaerobic Digestion Dynamics

**DOI:** 10.1101/2024.07.16.603753

**Authors:** Parsa Ghadermazi, Jorge L. Rico, Ethan Rimmelman, Siri Stenmark, Kenneth F. Reardon, Susan De Long, Siu Hung Joshua Chan

**Affiliations:** Department of Chemical and Biological Engineering, Colorado State University, 1301 Campus Delivery, Fort Collins, CO 80523, USA; Department of Civil and Environmental Engineering, Colorado State University, 1301 Campus Delivery, Fort Collins, CO 80523, USA

## Abstract

Handling the global food waste requires sustainable solutions. Anaerobic digestion (AD) is a common approach for waste remediation and bioenergy production which has been traditionally used mainly for methane production. Shifting the focus from methane to volatile fatty acids (VFAs), such as acetic, propionic, and butyric acids, offers promising alternatives due to their diverse applications and higher added value. Achieving desired AD product distributions requires a comprehensive understanding of factors like temperature, pH, feedstock composition, and the complex microbial dynamics inherent in AD. Various AD modeling approaches exist, from simple equations to complex flux balance analysis (FBA) and machine learning (ML) model. The Anaerobic Digestion Model No. 1 (ADM 1) is a commonly used kinetic model, striking a reasonable balance between parameter requirements and biochemical details involved in the model. Yet, it falls short in capturing specific VFAs like caproic acid and integrating microbial information directly. We present ADToolbox, a Python package for modeling AD metabolism. ADToolbox incorporates metagenomic information into an enhanced ADM model. The model accommodates a more detailed feedstock degradation model and VFA and methanogenesis metabolism. ADToolbox provides a variety of interfaces such as command line interface and an interactive web interface line interface, and a Python API, facilitating large-scale metagenomic analyses and modeling simulations. In this article we indicate that prioritizing the microbial aspect of AD enhances flexibility and predictive power in terms of VFA production accuracy, contributing to sustainable waste management strategies. Explore ADToolbox at https://chan-csu.github.io/ADToolbox for detailed documentation.

## Introduction

Every year, around 1.3 billion tons of food waste is produced worldwide, and the US alone is responsible for producing about 60 million tons this waste [1]. According to the United States Department of Agriculture, municipal solid waste landfill is the third largest source of methane, which is a greenhouse gas with 28 times larger global warming potential than carbon dioxide, emission [2].

Anaerobic digestion (AD) is a promising and widely used approach in bioremediation [3], [4], [5], organic waste valorization to biofuels [3], [6] and value-added chemicals [3], [7]. During anaerobic digestion, a complex network of microorganisms, breakdown the large molecules in the feedstock and obtain energy and precursors required for growth by metabolizing the resulting small molecules. As a result of such metabolism activities, products like methane, hydrogen, volatile fatty acids (VFAs), carbon dioxide are produced.

Methane production has traditionally been mainly the focus of AD [3], [8]. However, this pattern has been changing recently due to the low value of methane while still competitive resources for making methane exists. Additionally, methane is a greenhouse gas with high global warming potential which motivates for changing the AD process to products beyond methane [3], [9].

VFAs are interesting alternative products of AD. VFAs such as acetic, propionic, and butyric acids are essential chemical building blocks used extensively in the food and pharmaceutical industries, as well as in plastic production and wastewater treatment [10], [11]. Additionally, VFAs are strong candidates as substrates for bioenergy production [11].

Channeling the flow of metabolites from methane to VFAs requires thorough information about the influence of multiple factors on the outcome of AD. Temperature, pH, composition of feedstock have been shown to influence the AD product distribution significantly [12], [13], [14], [15]. More importantly AD is happening as result of the metabolism of a complex community of microbial cells, and the metabolism of the AD microbiome determines the distribution of the final products of the process [8], [16]. As a result, understanding the governing mechanisms in such communities is vital to engineering and optimization of AD. This matter becomes possible in the presence of models that describe AD outputs as a function of the inputs to the process.

Complexity of anaerobic digestion has fostered different approaches in modeling AD which compromise between multiple factors such as complexity of the model, the number of required parameters, explainability, and the generality of the model.

The simplest form of mathematical models for AD includes a single mathematical equation that relates one output to one or multiple input variables representing the characteristics of the feedstock and the inoculum source that is used in the process [17], [18], [19]. Although these models offer a simple way to model AD, they suffer from multiple problems such as lack of interpretability, lack of generality, and inability in predicting the systems dynamics [17].

A rich variety of mathematical models of AD use a single or a system of multiple differential equations to describe the dynamics of AD. A number of these variants use only a single ordinary differential equation, ODE, which usually relate the methane production rates at each state with one or more characteristics of the feed stock at that state [17], [20]. The main advantage of such models is the simplicity to use and that they usually do not need a wealth of experimental data for parameter tuning. However, such simple model will inevitably lose the flexibility and generality when applied to a different AD system.

On the other end of complexity spectrum lies microbial metabolic network models and flux balance analysis (FBA). FBA based approaches have been used in multiple studies related to AD [21], [22], [23]. FBA aims to predict the phenotype of a cell from the underlying metabolic model by assuming steady state condition across the cell and constructing a linear programming problem and solving for a given objective function such as biomass production of the cell [24]. FBA also can be extended to dynamic simulation of heterogenous microbial systems [25]. There are several inherent limitations in using FBA and DFBA, for example the instantaneous biomass maximization assumption that does not always describe observed microbial interactions [26], [27], [28]. Additionally, parameters like temperature and pH are shown to be critical in determining the performance of AD reactors which are often ignored in FBA-based approaches. Despite these limitations, FBA will continue to be an interesting venue for modeling AD as it provides a detailed model structure that links microbial growth and metabolism[29].

Machine learning (ML) models can capture the relationship between inputs and outputs of AD by learning directly from the experimental data provided to the model without the necessity of knowing the underlying kinetic equations governing the biochemical reactions of the system. Although multiple studies have shown the predictive power of ML algorithms in AD or similar environments [30], [31], [32], [33], [34], their generality, given the complexity of the underlying microbiome is questionable, especially, in the absence of plenty of data for training the models which is usually the case in AD studies [35], [36]. To incorporate the high-dimensional metagenomics features, gene or taxa abundances, into ML models that aim to predict AD metabolism, one needs to find features that exist in a lower-dimensional space. For example, [37] converted the taxonomy data from 16s sequencing of AD microbiome at phylum-level to microbial features in the various ML models along with other factors that are known to affect AD to predict methane production. One limitation of this approach is that the metabolic capabilities of taxa in the same taxonomic group, phylum in this case, is assumed to be identical while this is shown to be an inaccurate assumption (See “Results: *<u>A large-scale amplicon study”</u>*). Choosing more low-level taxonomic features can alleviate this issue to some extent, such as species level features. However, this will increase the number of required features that can be use across different AD systems which in turn requires more experimental data for reasonable accuracy. Additionally, inferring mechanisms behind the model prediction in ML models is not straightforward if not impossible.

Alternative to the aforementioned approaches, a family of simplified kinetic models try to strike a balance between the number of required parameters and the level of microbial/biochemical details included. Anaerobic Digestion Model #1 (ADM1), which was introduced in 2002 [38], is probably the most widely used model in this group. This model groups multiple biochemical reactions into a larger class of ADM reactions and these reactions follow a determined kinetics. In this way the model is limited to 28 reactions and 38 compounds. Using ADM1 reactions, Chemical Oxygen Demand (COD) balance across the bioreactor of interest results in a system of ordinary differential equations which is solved numerically to simulate the concentration profiles against time.

While providing useful information and simulations for the AD process, there are two aspects of insufficiency of ADM1 that need to be addressed for optimizing VFA production from AD. First, ADM1 has been traditionally used for modeling methane production from different feedstocks. With more attention shifted to VFA production, new AD models should include more details about VFA metabolism. For example, caproic acid production is observed in many anaerobic digestion bioreactors [16], [39], [40], [41]. and electron donors such as lactate and ethanol are found to be important for chain elongation during this process [42], [43], [44], [45], [46]. However, they are commonly missing from ADM1 and its derivatives. Second, metagenomics-derived microbial information about the samples cannot be used directly in the ADM model upfront. New AD models that take advantage of the wealth of information from metagenomics data for AD are likely to benefit the capability of making predictions.

This research aims to address the challenges associated with modeling the AD process by prioritizing the fact that anaerobic digestion is a microbial process, and the AD microbiome genotype is used as a direct input to an AD model to ensure flexibility and predictive power. Here we introduce ADToolbox as a set of computational tools bundled as a Python package. ADToolbox includes an enhanced ADM kinetic model that integrates metagenomics information from AD samples and adds ethanol, lactate, caproic acid components as state variables. The enhanced ADM model we are proposing also incorporates the concept of total suspended vs. dissolved solids for a more complex feedstock degradation, which is inspired from [47]. ADToolbox offers a command line interface, an interactive web application that could be easily hosted online out-of-the-box, and it also offers a Python API which is designed to handle large-scale metagenomics analyses and modeling simulations efficiently. ADToolbox is freely available at: https://github.com/chan-csu/ADToolbox. A detailed documentation for using ADToolbox is provided at: https://chan-csu.github.io/ADToolbox.

## Methods

An overall view of ADToolbox capabilities is provided in Figure 1. ADToolbox offers novel capabilities for modeling AD at different levels and it can be viewed as a combination of inter-connected modules. These modules work in harmony to make prediction of AD dynamics possible from microbiome data.

**Figure 1.**
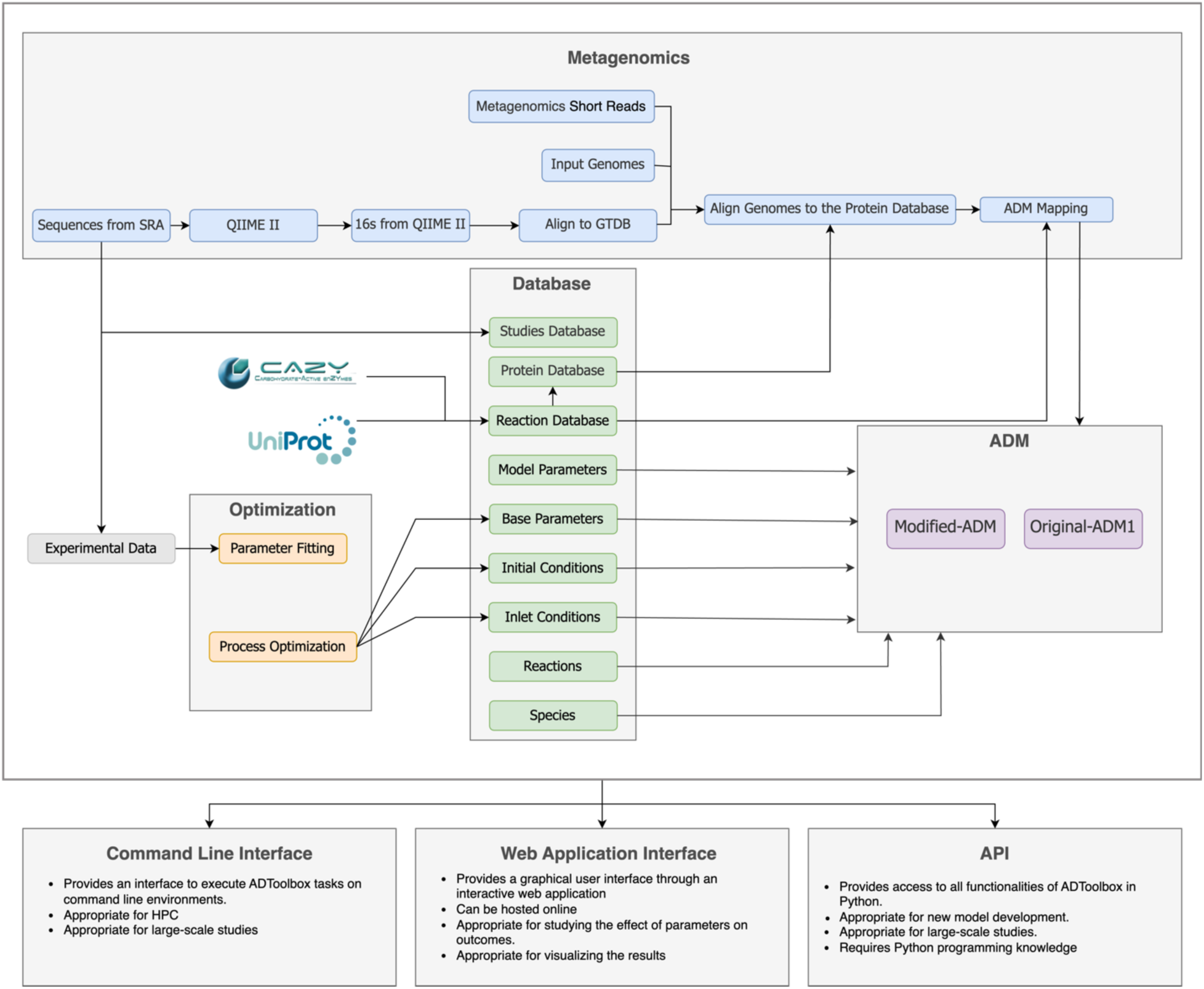
Overall view of relationship between ADToolbox functionalities. The goal of this package is to offer a parameterized model like ADM1 that can simplify and incorporate metagenomics data and offer a database of published studies to tune the model parameters on.

### ADM module

This module comes with two implementations of ADM and it is easily expandable to more ADM models. The first implementation is a rearrangement of the PyADM1 [48] implementation of the exact same original ADM1. ADToolbox offers a new implementation of an extended ADM model (called ‘eADM’ hereafter) that improves the ADM1 model in 1) Feedstock degradation: To take differentiate different feedstock with different degree of availability of the ingredients 2) Ethanol, Lactate, and Caproate metabolism: To allow chain-elongation functionality of the microbiomes 3) Methanogenesis is possible either from CO2 and H2 or from Acetate independently: To differentiate acetotrophic methanogenesis from hydrogenotrophic methanogenesis. The eADM model includes 50 reactions and 49 compounds. The model simulates a well-mixed anaerobic digestion bioreactor, and the dynamics of the system is described through a system of ordinary differential equations, ODEs. SciPy [49] is used for solving the system of ODEs.

Another important aspect of contributions other than the model kinetics is the user interface and visualization. The solution of this system can be either outputted in a tab separated file or visualized as an interactive web interface using a dash web application [50]. Users can interactively see the effect of any model parameters on the system dynamics. Additionally, this web application can show how flux is distributed through the metabolism network, an Escher map [51], at each timepoint in the simulation.

#### Hydrolysis

The reactions in this section simulate the hydrolysis step in AD which is responsible for breaking down the large molecules into smaller molecules that can be taken up by the cells and metabolized further. At first, the degradable part of the solid feedstock is comprised of two portions: Total suspended solids, TSS, and total dissolved solids, TDS. Degradation of each portion follows same linear kinetics as defined in ADM1. However, the kinetic parameters for degradation of each portion is different to reflect the different physicochemical rules governing each portion while keeping the simplicity of the model. This separation of faster vs. slower degradation kinetics was reported to be more predictive of the hydrolysis step [47]. In this step, TDS degrades into carbohydrates, lipids, proteins, and *soluble inerts*. On the other hand, TSS degrades into carbohydrates, lipids, proteins, and *particulate inerts*. In turn, carbohydrates are converted into sugars, proteins into amino acids, and lipids into fatty acids (Figure 2-A).

**Figure 2.**
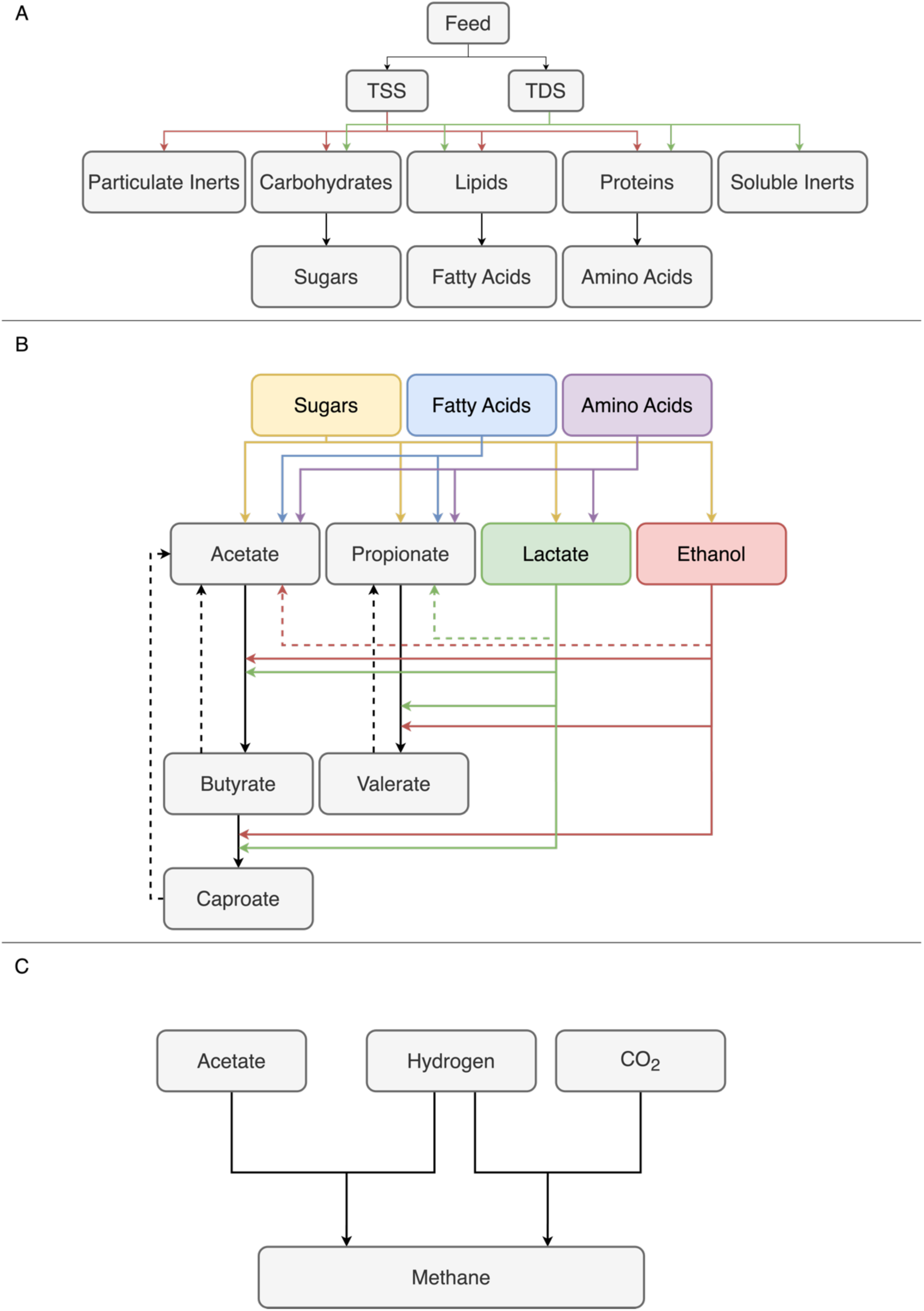
A high-level view of the modifications from ADM1 model A) This section shows that the soluble and insoluble part of the feedstock degrades at different rate. B) This section shows how acids are generated from sugars, proteins, and fatty acids. The dashed lines show the reactions in the reverse direction, oxidation reactions. C) Methanogenesis is now happening by either acetotrohphic methanogenesis and hydrogenotrophic methanogenesis.

**Figure 3.**
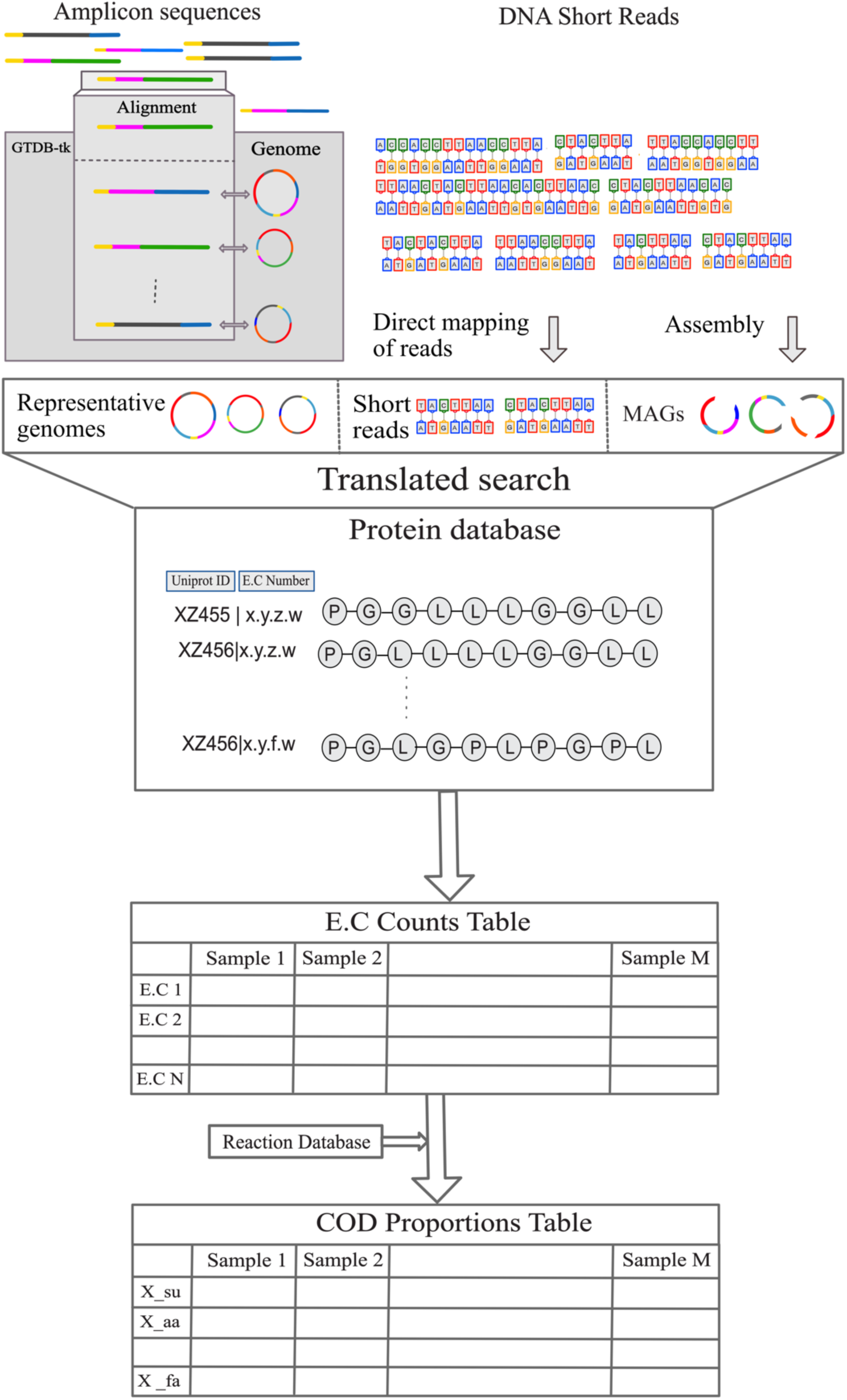
Metagenomics module of ADToolbox. This module can take metagenomics data, either 16s, Mags, or short reads, and aligns it with ADToolbox protein database. Finally, using the reaction database of ADToolbox, COD proportion of each microbial species in the modified ADM is estimated.

#### Acidogenesis and Acetogenesis

Small molecules resulted from the hydrolysis step are converted to acetate, lactate, ethanol, propionate, valerate and butyrate (Figure 2-B). All the expected VFAs are produced at this step. Butyrate is produced from elongating acetate and taking electrons from either ethanol or lactate. Similar mechanism elongates valerate from propionate and caproate from butyrate.

#### Methanogenesis

There are two methanogenesis pathways in eADM. First, similar to original ADM1, methane can be produced from acetate. Second, eADM implements methane production from hydrogen and carbon dioxide to decouple this mode of methanogenesis from acetotrophic methanogenesis (Figure 2-C). Generation of methane from CO_2_ and H_2_ is possible using the Wood-Ljungdahl pathway. This pathway also produces alcohols and fatty acids [52] from CO_2_ and H_2_. However, the suggested model does not cover this aspect of the metabolism yet.

### Database Module

This module handles any form of data that is required for utilizing ADToolbox functionalities. At the heart of this module is the ADToolbox reaction database. This database is a flat file that relates enzyme commission (EC) numbers for biochemical reactions to the higher-level ‘lumped’ reactions in eADM. The database is curated manually to include biochemical reactions relevant to the primary substrates and products in AD. As an example, EC 4.2.1.11, which represents phosphopyruvate hydratase, is mapped to the sugar uptake reaction (“uptake_of_sugars”) in eADM. This eADM reaction converts “sugars” in the system to acetate, propionate, ethanol, and lactate. This database includes 840 EC numbers that are mapped to ∼50 reactions in eADM.

Another database is ADToolbox’s protein database. This database is constructed from all the EC numbers in the reaction database by searching Uniprots for any protein sequence known to link to the query EC number. The resulting protein database contains about 35000 protein sequences which can be used to align short/long sequences of DNA or protein of interest for downstream qualitative/quantitative analysis.

ADToolbox also comes with a feed database that includes pre-processed data from literature about composition of different feedstock along with links to the source. Any item in this database could be used as an input to eADM.

Another important aspect of the database module is the “Studies” database. This database includes information about published datasets that can be used by ADToolbox. Any entry in this database includes all the information that is published for one experiment run: operating conditions, initial concentrations of the system and time-series data for concentration of compounds in the system.

ADToolbox’s “Studies” database also includes a table for storing and querying sequencing data available on the SRA repository. The stored studies could be either whole genome sequencing data or 16s amplicon sequencing data. The main goal of this table is to store different studies in a way to make them comparable regardless of the timepoints at which the experimental data are reported or the type of sequencing data that is used for the study of interest.

Any instance of the ADM model requires a set of parameters for simulations: kinetic parameters and operational parameters for the bioreactor of interest. These parameters are also included in the database.

More information about the databases can be found in: https://chan-csu.github.io/ADToolbox/API/#7-database

### Metagenomics Module

The goal of this module is to extract AD-related functionalities from metagenomics data. This data will later inform the modified-ADM model about the metabolic potentials of the samples used to study. This module provides several metagenomics functionalities that mostly rely on external software packages (see Figure 1). This module allows downloading sequences from SRA using a wrapper around SRA-Toolkit, processes 16s sequences using a QiimeII [53] wrapper, shotgun short reads and genome alignment using a wrapper around MMseqs2 [54] and a few functions for processing the outputs of these external tools. The final goal of this module is to convert microbial information, either 16s or shotgun metagenomics, into ADM model inputs. If 16s sequencing data is provided, first relative abundance of each taxon is obtained by using the wrapper function for QIIMEII. In the next step, representative genomes are found, if exists, for each taxon using VSEARCH [55] and GTDB-tk database [56]. Consequently, the representative genomes are aligned to the protein database using MMseqs2 to find the EC numbers that are found within the genome of each taxon. Finally, the abundance of a biochemical reaction is estimated by counting the presence/absence of the corresponding EC number of each genome scaled by the relative abundance of the corresponding taxon. On the other hand, if shotgun short reads are provided, EC number counts are directly calculated by mapping the reads to the protein database and then scaling by the method of interest such as total sum scaling. In either case, the scaled EC count table is transformed into the relative COD relation between microbial species in eADM. The final COD table includes relative abundance of each microbial group in eADM. To calculate the absolute value of initial concentrations of each of these microbial species, total amount of the inoculum, in COD is multiplied by the relative abundance of that taxon and finally divided by the liquid volume of the reactor. This method approximates the initial concentrations of the microbial groups in a way that is compatible to eADM and provides a way to incorporate metagenomics data into ADM-like models.

### Optimization Module

This module is responsible for: 1) calibrating the model parameters using the experimentally observed data, and 2) optimizing process conditions for shifting the product profile towards our engineering goal. The non-linearity and instability inherent to kinetic models like ADM [57] favors metaheuristic optimization approaches. Accordingly, three different optimization approaches have been implemented in ADToolbox to achieve these two goals. Although three different approaches are offered in the optimization module of ADToolbox, but they all provide the same simple interface to optimize the models.

The first method relies on Bayesian Blackbox optimization method where the objective function is approximated with a function that fits the objective can be used to guide the search for the global optima. OpenBox library [58] in python is used as the implementation for Bayesian Blackbox optimization approach.

Genetic Algorithm, GA, is another approach that has been used with success for parameter fitting in kinetic models [59]. GA is employed in ADToolbox using pyGAD library [60].

The third optimization approach is an in-house algorithm and is conceptually like Bayesian Blackbox method. We developed this algorithm to improve the flexibility of function approximation in existing implementations of Bayesian Blackbox approach. In this method, first an initial population of the objective function value is calculated for different set of parameters. Next, a randomly weighted neural network estimates the objective values for the points that are already calculated and then using gradient decent algorithm the network parameters are changed to fit the actual objective values as good as possible. Afterwards, the network parameters are fixed, and gradient descent is applied to the network inputs, ADM parameters, to minimize the objective value. This step suggests a parameter tensor that could be used in the ADM model to calculate the new objective values. The new objective values are in turn used to train the neural network to estimate the objective function better in an iterative process. To avoid sticking in local optima, when the suggested parameters don’t change any longer, a new solution is suggested by drawing from a normal distribution around the current local optima to balance exploration with exploitation. Based on our analyses, this method proved to be more effective in calibrating model parameters then the former approaches mentioned, and we have done all of the optimizations mentioned later using this approach. The neural networks are defined and trained using PyTorch [61] library.

One attractive characteristic of the optimization module is that it connects easily with the studies database, and any combination of experimental data in the database can be queried for parameter calibration or model validation. Examples on how to use this module is provided at: https://chan-csu.github.io/ADToolbox/parameter_tuning.html

### Workflow

As ADM1 provides a good balance between the number of model parameters and the biochemical detail of the core model, the modified version of ADM, eADM, that additionally takes lactate, ethanol, caproate into account is at the heart of ADToolbox. Most of such reactions involve a microbial group that catalyzes the corresponding reaction. We used this characteristic of the family of ADM models to integrate metagenomics data into the model. Each microbial group in ADM is linked to one or more EC numbers in the manually curated reaction database of ADToolbox. Each EC number is also linked to as many prokaryotic or archaeal proteins found on Uniprot through ADToolbox’s protein database. As a result of this computational capability, shotgun metagenomics short reads, metagenome assembled genomes (MAGs), or amplicon sequencing data can be used to estimate the relative abundance of each group of microbial groups in the modified ADM model. Amplicon sequencing data can be processed either previously by the users or through the QIIMEII functionality provided with ADToolbox. The outputs of QIIMEII pipeline is then further processed to fetch the representative genomes for each taxa if possible. Representative sequence, repseqs, of each OTU, are aligned to GTDB-tk database to find the representative genome identifiers. Consequently, the genomes are downloaded from NCBI server. Although this method allows for flow of information from amplicon sequencing data, but the outputs of the pipeline should be treated more carefully compared to shotgun metagenomics short reads or MAGs as loss of information is inevitable when inferring function purely from amplicon sequencing data, see “Results: *<u>A large-scale amplicon study”</u>*

ADToolbox is designed to work with computing clusters and is capable of handling large metagenomics data efficiently. The package is designed to work with Docker and Singularity to maximize reproducibility and portability [62], [63].

### Implementation

ADToolbox is implemented as an open-source Python package. All the source codes are available through the GitHub repository for this project: https://github.com/chan-csu/ADToolbox

A detailed documentation website also exists that guides the users through the functionalities of ADToolbox along with the case studies that are mentioned in this paper.

## Results

The main novelty of ADToolbox is that it integrates genetic information in the form of DNA sequencing data from the environment of interest with an extended ADM model (eADM) to provide a mechanistic view about the environment of interest. In this section we show how these functionalities can be used to predict the dynamics of AD system across different microbiomes and how engineering strategies can be extracted from the simulations.

### Incorporating 16s amplicon sequencing data to predict VFA production

When expanding the original ADM1 model, we needed to include new kinetic parameters for the novel reactions in eADM. These parameters should be calibrated to make sure they can capture the kinetic rules governing anaerobic digestion. This necessitated using a study that includes microbiome data and time-resolved concentration data for at least the VFAs. The effect of inoculum source on VFA concentrations are explored in multiple studies [16], [64]. For calibrating the parameters of the model, we chose the study by [65]. This study explores the relationship between microbiome composition, in the form of 16s amplicon sequencing data, and VFA concentrations across three different microbiomes: Anaerobic sludge (AS), thickened waste anaerobic sludge (TWAS), and a mixture of AS and TWAS (TWAS-AS).

After converting the 16s data into eADM’s microbial groups relative abundance for samples in the beginning of the experiment, we sought to check if the different microbiomes are compositionally different in the space of eADM’s microbial groups. Figure 4-A shows the non-metric multidimensional scaling (NMDS) plot with Bray-Curtis distance between the samples. It is evident from this figure that the metagenomics pipeline that converts metagenomics data to relative abundance of ADM microbial groups can differentiate the samples from different microbiomes.

**Figure 4.**
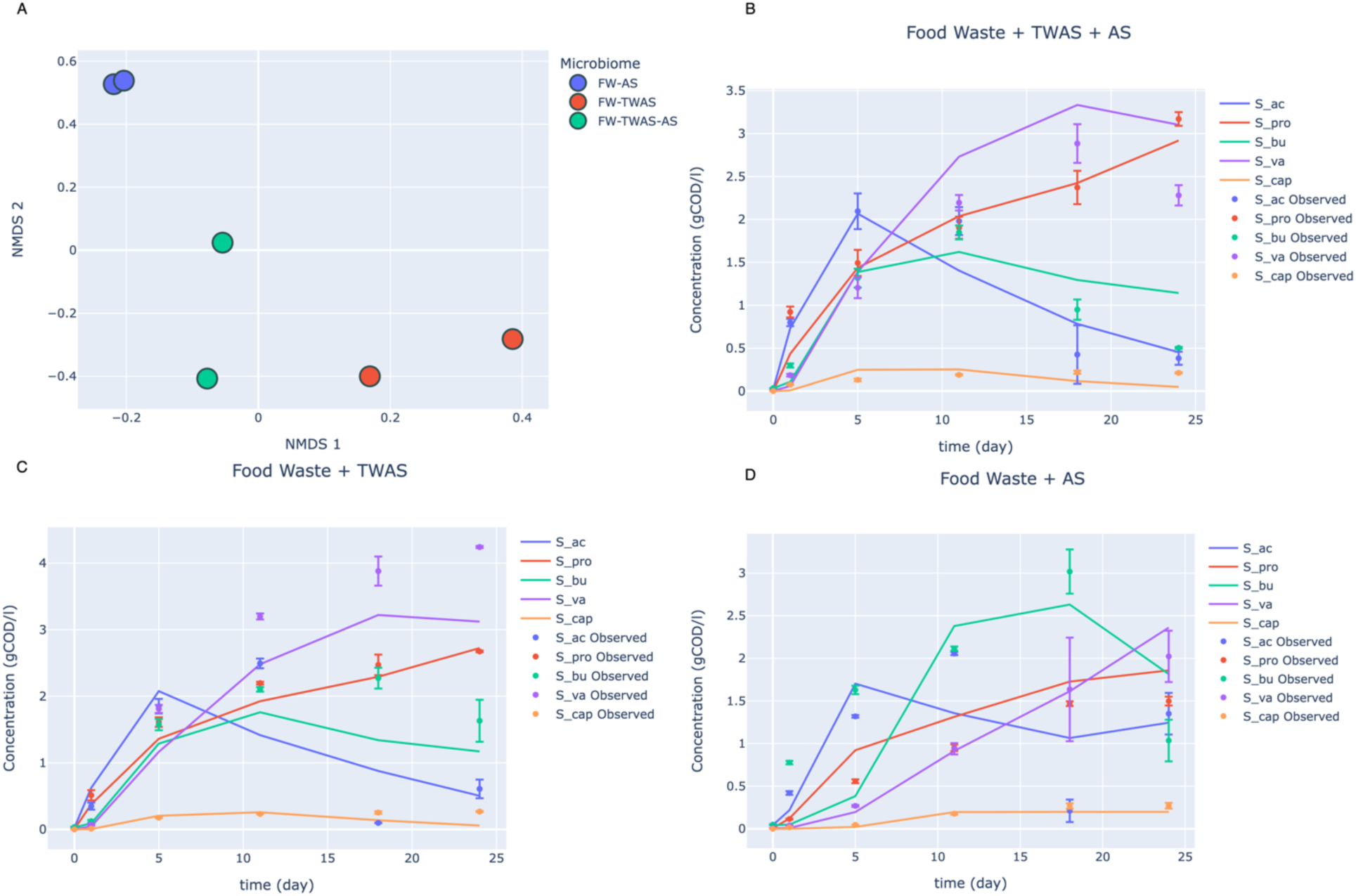
Performance of ADToolbox in prediction AD concentration profiles over 3 different microbiomes. A) Results of the metagenomics pipeline: NMDS plot showing the relationship between the samples when 16s data is mapped to modified-ADM’s microbial groups. B, C, and D) VFA concentration profiles for model prediction compared to the experimental data for three different microbiomes on food waste: TWAS+AS, TWAS, and AS respectively. The error bars show the standard error for each microbiome across different replicates. The experimental data is from [64]

Figure 4 B-D describes the model prediction compared to the experimental data reported in [64]. Overall, the model can describe the VFA production dynamics of AD across three microbiomes reasonably well for all data points (r^2^=0.85). It is worth mentioning that the only factor that is different across figures 4 B-D is the inoculum source.

Same simulation results indicate that using metagenomics data to inform e-ADM model helps with more accurately predicting methane production differences between different microbiomes (Supplementary Figure 2). It is evident that both order and behavior of methane production curves closely resembles the experimental data for methane concentration in [65].

The studies mentioned here are all included in the “studies” database of ADToolbox and they provide a ground for fairly comparing different studies on AD systems in a way that is compatible with the structure of ADM-like models.

### A large-scale amplicon study

Incorporation of metagenomics data for quantitative predictions is a useful idea because it allows us to examine the performance of different microbiomes in AD environment and minimize expensive and time-consuming experiments through model-guided experiment design.

One question that is relevant to this concept is: can we define a core microbiome for the hosts at function level? If the answer to this question is yes, then we can select a host that performs better than others and use it to inoculate the AD reactor of interest. To answer this question, and prove the scalability of ADToolbox with large datasets, we analyzed 5700 16s samples fetched from the sequence read archive, SRA, belonging to the gut microbiome of animals with potential application in AD process. Processing this many samples in a short time, less than 24h, became possible because ADToolbox is designed to work with clusters efficiently. Figure 5-A shows t-distributed Stochastic Neighbor Embedding (TSNE) plot of taxonomy of all the samples used in this study. At this level we observed clustering between samples belonging to the same host when individual taxa were used as features. However, mapping this compositional data into functional space did not reveal the same clustering for the samples. There could be different reasons behind this observation. We think an important reason is the information loss when converting 16s sequencing data to functional information since this approach is relying on representative genomes only instead of actual functional genes in the sample of interest. This issue becomes even more pronounced when analyzing samples across multiple studies as more variability is introduced to the input data.

**Figure 5.**
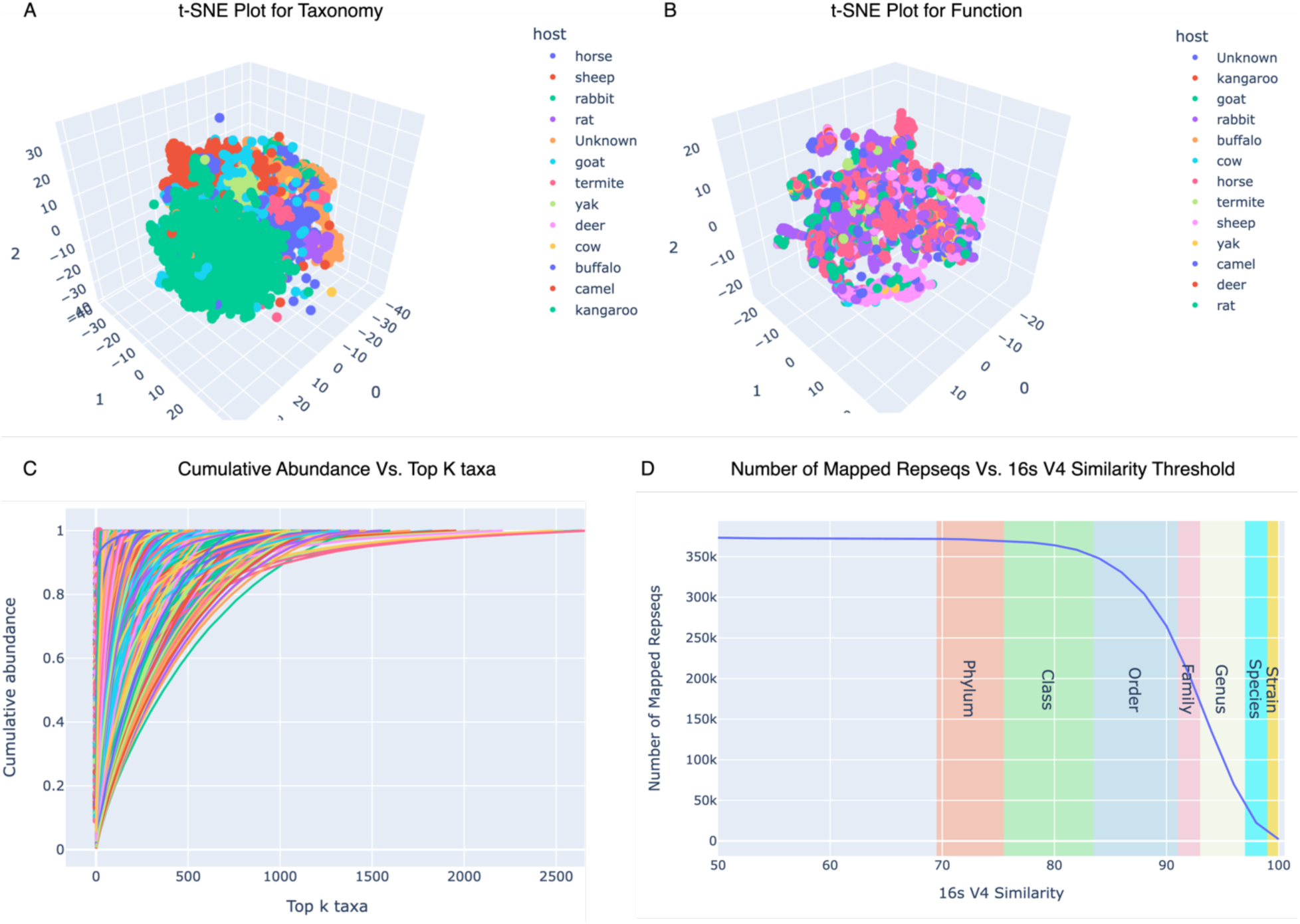
Limitations of functional interpretation of 16s data. A) t-SNE plot for taxonomy data shows clustering for samples that belong to the same host. B) t-SNE plot for the same data mapped to functional space shows no relationship between samples coming from the same host. C) This plot shows the complexity of the 5700 samples that are used in this study. For some samples even top 1000 taxa only cover 80% of the community. D) This plot shows the trade-off that is inherent to function inference from amplicon sequencing data. Higher threshold causes lower number of mapped genomes for the taxa present in the samples while the genomes represent the taxa more closely than a lower threshold value.

Figure 5 explains the limitations of using 16s sequencing data as functional information that can fed to the ADM model.

Shotgun metagenomics on the other hand, provides a direct approach to infer functions from samples and we hypothesized that using shotgun metagenomics data gives the model more predictive power which has been explored in the next section.

### A shotgun metagenomics study

Motivated by the more functional information available in shotgun metagenomics data and the increasing number of such sequencing studies, we extended the suggested pipeline for incorporating shotgun sequencing data. Given metagenomics short reads there are two separate routes that one can take: Assembling the reads to produce metagenome-assembled genomes, MAGs, or directly map the short reads to a reference database. ADToolbox does not provide assembly functionalities. However, if MAGs and their corresponding relative abundance is provided, then this information can smoothly be incorporated similar to the method explained for 16s sequencing data but in this case the assembled genomes are used instead of reference genomes. On the other hand, using ADToolbox, short reads can be directly aligned to the curated protein database of ADToolbox and then translated into relative abundance of the microbial groups in the eADM model and we selected this approach for this study.

Among many shotgun metagenomics candidates available we selected a study done by [66]. Although the goal of this study was not to compare VFA production in AD reactors but the rich metagenomics data on the candidate animals made it a suitable candidate to explore the potentials of these microbiomes in AD system. The dataset for this study contained 12 metagenomics samples, 3 samples for cow, 3 samples for red deer, 2 samples for sheep, and 2 samples for reindeer. The typical diet of these animals makes them interesting candidate for anaerobic digestion as their diet usually includes large polymers commonly used in AD bioreactors such as cellulose.

After mapping the short reads of the samples to the protein database of ADToolbox and extracting E.C numbers, the relative COD distribution across microbial groups of the eADM model were calculated as explained in the method section. Similar to the previous case study, we constructed NMDS plot for relative abundance of eADM microbial groups (Figure 6). It can be observed from Figure 6-A that the samples of different hosts are separated in this NMDS plot. This observation suggests that the inputs to the modified ADM model is different for cow and goat samples, and as a result, we can explore how this difference in the microbiomes translate into differences in VFA production profiles. To compare the potentials of these microbiomes in VFA production, we simulated the VFA concentrations over time on all 12 samples mentioned above and have summarized the results in Figure 6B-G. The feedstock that was used for this study was selected to emulate cellulosic media with total COD concentration of 20 gCOD/l with 100% insoluble carbohydrates, 100% TSS.

**Figure 6.**
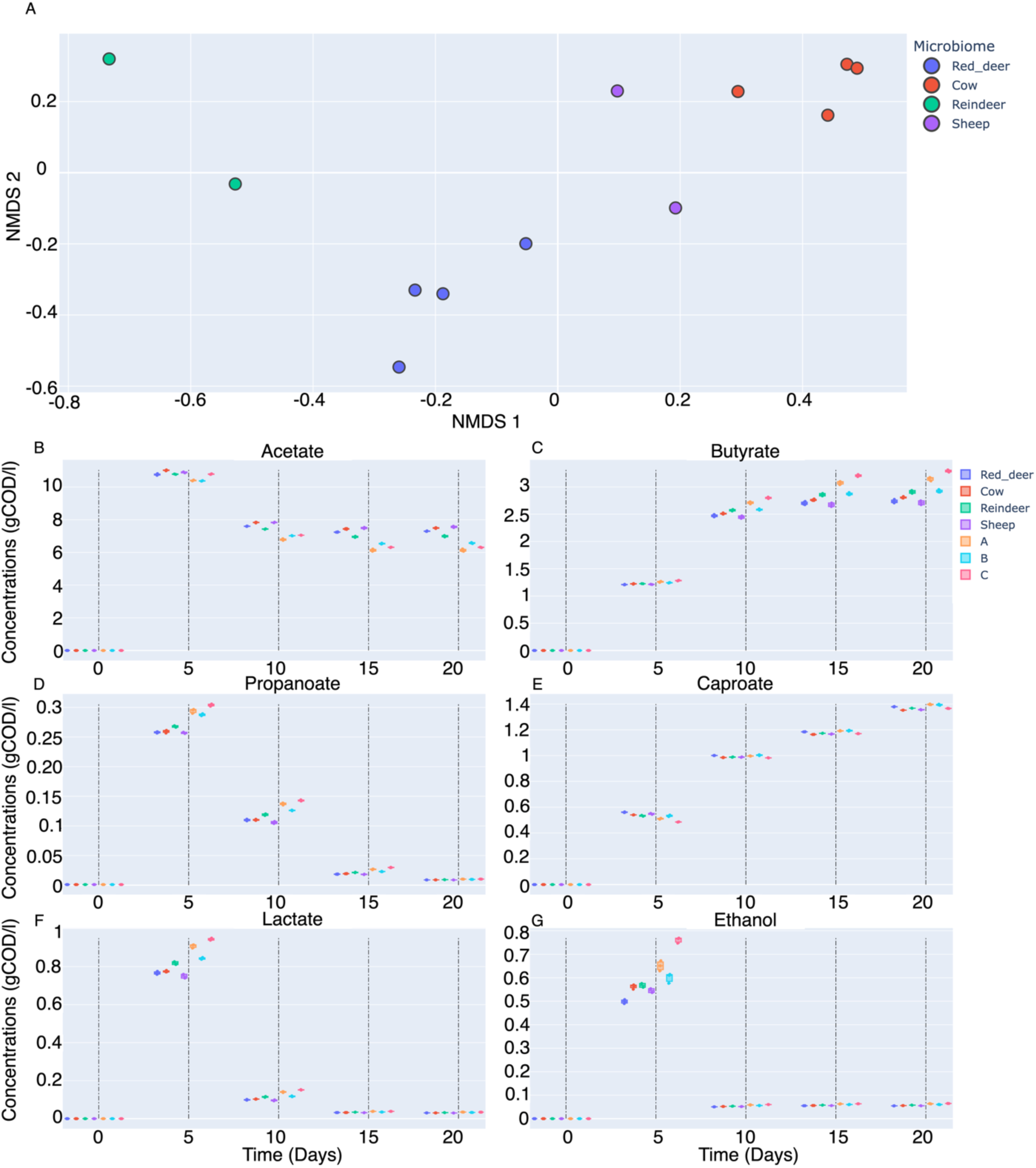
Incorporating information from metagenomics short reads into the modified ADM model. A) NMDS plot describing the similarities and differences in the relative abundance of microbial groups in the modified ADM model inferred from the shotgun metagenomics data from ref. [66]. B-G) Simulated concentration profiles for different compounds over 20 days in an anaerobic digestion reactor. Colors are representing the corresponding microbiome. For comparison purpose, we have included the day-0 samples from ref. [67] to the right of the dashed lines at each time point. Note that A, B, and C follow the same naming convention as described in [67] where they correspond to 16s amplicon sequencing data from anaerobic sludge, bison rumen, and cow rumen respectively.

Interestingly, the suggested workflow can provide a common ground to compare simulation results that are based on different sequencing technologies because regardless of the technology, sequencing data is mapped to a low dimensional space that is represented by the relative abundance of eADM’s microbial groups. For comparison purpose, we have additionally included the 16s sequencing data from [16]. We can see that depending on the microbiome composition the simulated VFA profiles are different. This difference is solely because of incorporating the sequencing data from these environments are included in the model. All the other inputs to the model are similar across the simulations.

Interestingly, during the process of converting the high-dimensional functional representation of the metagenomics data to the lower-dimensional representation of modified ADM microbial groups preserves the separation between the different microbiomes, Figure 6-A. The difference in the microbial groups abundances results in the difference in concentration profiles for each microbiome.

Comparing the simulation results based on the shotgun metagenomics samples from[66] did not match the pattern for VFA production that was reported in [16]. As an example, [16] reported that cow produces relatively high concentration of propionic acid while the simulations for the cow samples from [66] shows low propionic acid production compared to other microbiomes, Figure 6-D (left of the dashed line). For this reason, we decided to also run similar simulations side by side using the 16s data from [16]. Using the 16s data ADToolbox was able to predict the highest propionic acid production for cow microbiome using the sequencing data from [16], Figure 6-D (right side of the dashed lines). We attribute this discrepancy between different studies to the expected difference in the microbiomes of similar hosts that reside in different environments, cows in this case. To some extent, we were expecting this behavior as it was similar to the observations in Figure 5: Microbiomes of the same animals were vastly different between different studies in the functional space. The simulations were also consistent for other VFAs which provides further evidence that integrating metagenomics information into the ADM-like models can increase their predictive power while keeping a mechanistic view of the process.

One special capability of ADToolbox is that for each of these simulations, an interactive Escher map [51] describing the metabolism of the microbiome at each instant of time can be generated which provides a more mechanistic overview to the metabolic flux distribution across the network. For more information and examples, the readers are advised to visit the documentation website for ADToolbox: https://chan-csu.github.io/ADToolbox/

## Discussion

Anaerobic digestion is a process that is carried out by microorganisms dwelling in anaerobic condition. However, most of the modelling/engineering effort in this field treat the microbiome as a black box and Information flow from genotype to quantitatively describing the microbiome phenotype is often ignored in AD research. Multiple published work [68], [69], [70], [71] use microbiome data through approaches like FBA, but the complexity of AD does not allow for extensive parametrization in these approaches. As an example, to the extent of our knowledge, there is no study that explores the effect of different inoculum sources on AD dynamics that can incorporate the effect of critical parameters such as pH and temperature. In this study we have made this possible by integrating metagenomics data, both amplicon sequencing and shotgun metagenomics data, into a modified version of the ADM1 model. In this approach the sequencing data is first aligned to the protein database. Next the aligned reads are then mapped to the microbial groups that are represented in ADM-like models. This step reduces the dimension of the data from a high-dimensional gene feature space into lower dimensional “microbial group” features. In the case of our modified ADM there are 18 different microbial groups. This means that any type of sequencing data from a sample is finally represented by 18 relative abundance values. This empowers the ADToolbox in many ways, including:

- The number of microbial features is still low, and this limits the number of the parameters that the model requires to predict AD performance.
- Assigns high-level roles to individual taxa based on the low-level gene features. For example, based on the alignment results and the mapping to the ADM model this workflow can specify the genes that a specific taxon contains for hydrolysis step of AD.
- Provides a unifying ground where studies that have used different types of sequencing techniques can be combined. This is possible because the sequencing data of any kind is finally converted to the relative abundance of the microbial groups in the ADM model, Figure 6 B-G.

We think that the last point is a promising capability because it enables us to build a database of published studies and used them for calibrating the model parameters regardless of the sequencing technology that is used in that study. This makes the model more generalizable and increases predictive power. As an example, the two studies that are mentioned in the results section are now included in the studies database of ADToolbox, and any combination of the experiments can be seamlessly used to calibrate or validate the model. See the documentation website for more information.

We simulated multiple studies that provided microbiome data and our results showed an interesting pattern. While the microbiome differences were reflected in the model outputs, we were not able to define a core function for the same host, Figure 5-B. We also saw the same pattern when we compared the cow samples between two studies in Figure 6 B-G. A possible explanation for this observation is that there are many factors influencing the microbiome composition and function such as location, diet, etc. which could be different between the hosts.

Although any kind of sequencing data can be used as input to the pipeline, when we scaled up to a large number of publicly available data for 16s amplicon sequencing data, we found that inferring function merely based on 16s data could have potential issues and we explained two reasons in Figure 5 C-D. Based on these results we recommend checking what percentage of taxa in the samples are covered by available representative genomes. Since with shotgun metagenomics data the information about the genes in a sample can be directly retrieved, we created a pipeline to directly map the reads to the ADToolbox protein database. This method is independent of representative genomes, and as a result, will produce a more realistic view of the function of the microbiome. Results from mapping short reads of shotgun metagenomics data also revealed that the functional differences between metagenomics samples preserves after mapping high dimensional representation of the metagenomics data in terms of EC numbers are mapped to ADM microbial group’s abundances. This is a promising finding that the metagenomics data can be used to differentiate the effect of different inoculum sources before running any experiments, regardless of the type of sequencing data. However, with dropping prices of shotgun sequencing, such data can be used to model the AD dynamics more accurately. Another observation from the simulations shown in Figure 6 is that the most prominent difference in concentration profiles belongs to lactate and ethanol. Although, the model parameters have not been calibrated using lactate or ethanol concentrations, the difference in VFA profiles observed for different microbiota is attributed to their capabilities in producing electron doners in the form of ethanol or lactate because of the underlying biochemistry of the model (Figure 2-B).

One clear advantage of using metagenomics data in ADM-like models is that it still has a mechanistic view of AD metabolism. Supplementary Figure 1 shows an instance of the web interface offered by ADToolbox where at each timepoint we can see how metabolic flux is distributed across the metabolic network. This information can help us understand what the main bottlenecks of the process at each timepoint are and what might be the potential intervention strategies that can improve the efficiency of the process.

We improved the kinetic model of ADM1 [72] to account for the changes in the concentration of important compounds and addressed a number of limitations in the underlying kinetics for a few of the reactions [57]. The main improvement was on VFA production. Although ADM1 has been widely used for modeling AD, the main focus of ADM1 is on methane production. Recently more studies have shifted their attention towards VFA production [16], [64]. This shift of attention motivated developing an extended version of ADM that include more information about VFA production which resulted in including caproic acid, ethanol, and lactate into a modified version of ADM1. To distinguish between the kinetics of degrading soluble and insoluble portions of the feedstock, we divide the feedstock to total suspended solids and total dissolved solids with similar linear kinetics but with a separate kinetic parameter. Finally, we separated methanogenesis to two separate terms to distinguish acetotrophic methanogenesis and hydrogenotrophic methanogenesis, Figure 2.

Given the complexity of AD system, a simplistic model like ADM could fail to predict the nuances in AD dynamics. However, we claim that our pipeline can go beyond the ADM-like models towards more data driven approaches. As mentioned before, the metagenomics data is converted into a lower-dimensional representation. This provides the possibility to use these metagenomics feature in machine learning models. In this case the advantage of this method is that the number of features does not require large amount of data for training a data-driven model. Previous work on using machine learning models to incorporate metagenomics for predicting AD outputs usually did not have enough data to train a large model. To overcome this issue, usually the metagenomics data are mapped to higher taxonomic level such as phylum level. However, functional similarity between the members of such high taxonomic levels is questionable. Our approach on the other hand, relies on the genomic content of the constituent taxa of the system. For this reason, we think the metagenomic features from ADToolbox can more precisely describe the functional potential of the system in a low-dimensional space, and is a good future direction for building machine learning models for AD.

## Conclusion

ADToolbox is an opensource python package that is designed for efficiently modelling the AD process as a dynamic system. This package can incorporate metagenomics data into a kinetic model that is extended from ADM1. ADToolbox is designed with the goal of making quantitative prediction for AD system from metagenomics data and can further improve our understanding of AD metabolism.

## Acknowledgement

This work was supported by the US Department of Energy Office of Energy Efficiency and Renewable Energy, Bioenergy Technologies Office under award number DE-EE0008923 This work utilized the Alpine high performance computing resource at the University of Colorado Boulder. Alpine is jointly funded by the University of Colorado Boulder, the University of Colorado Anschutz, and Colorado State University.

