## Supplementary Figures for "ADToolbox: Incorporating Metagenomics Data for Improved Prediction of Anaerobic Digestion Dynamics"

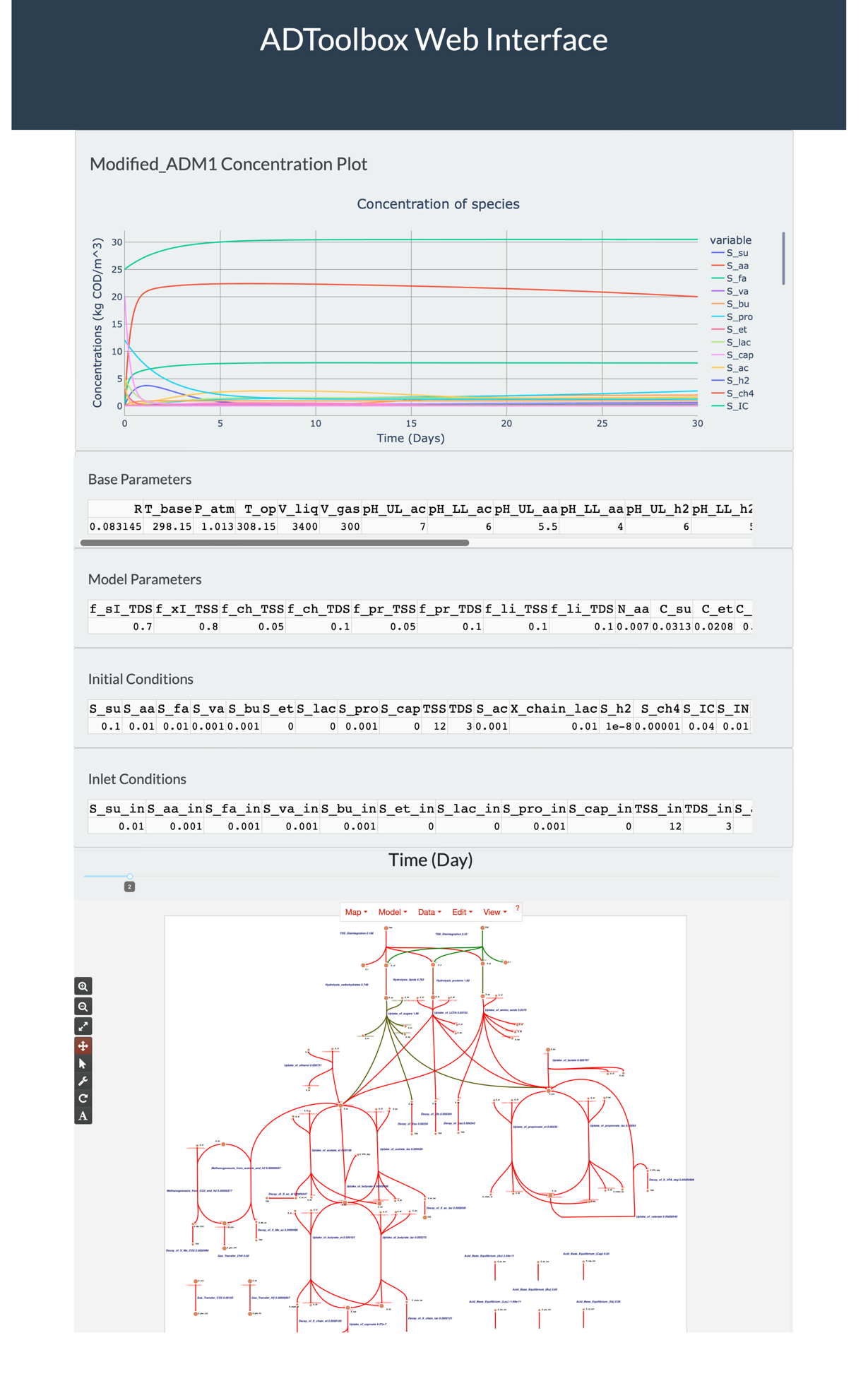

**Supplementary Figure 1** An example of ADToolbox web interface. This interface enables the user to see the real-time effect of the model parameters on concentration profiles as well as a time-resolved metabolism map for an example of ADM model (e-ADM in this case).

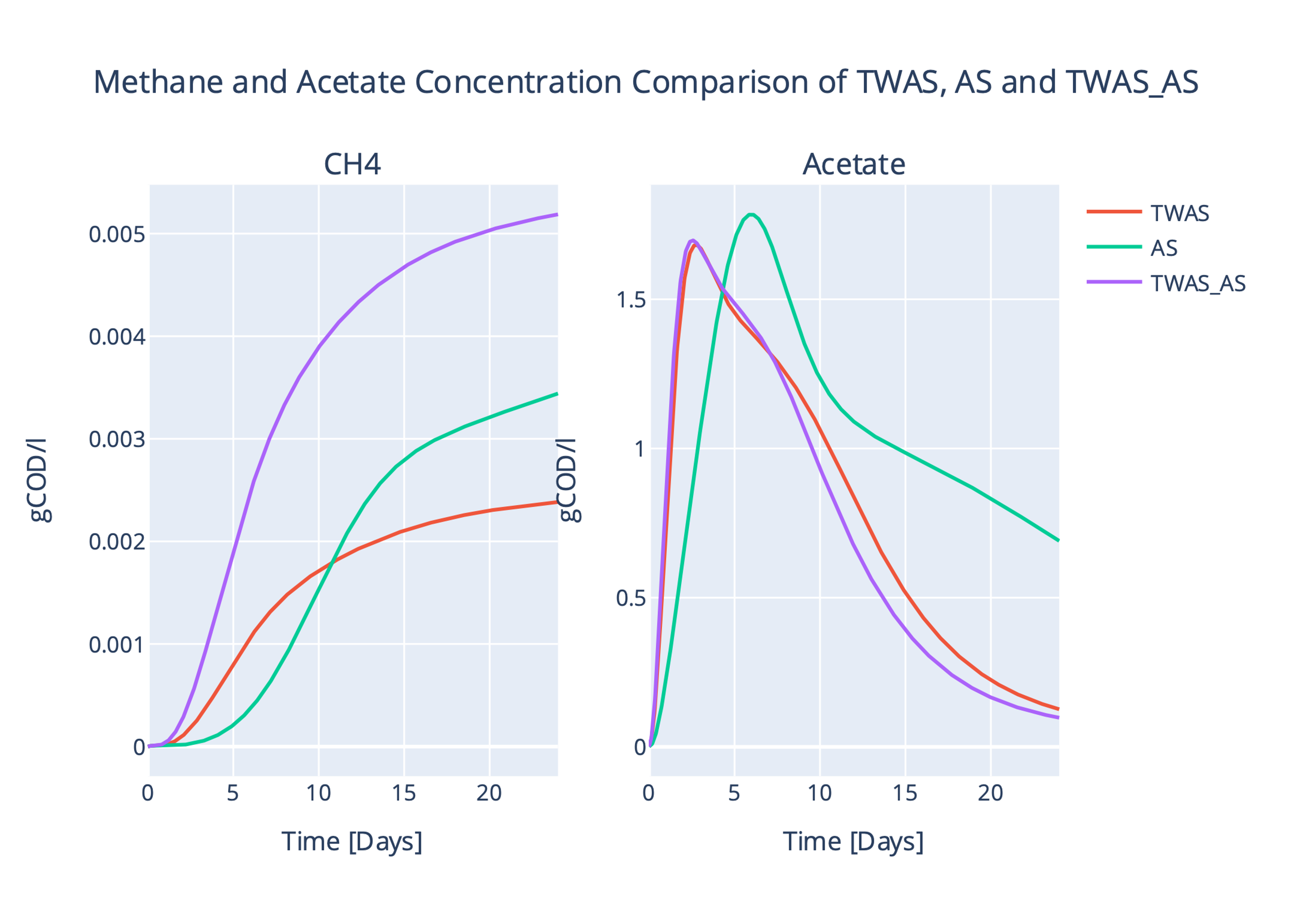

**Supplementary Figure 2** Methane and acetate concentration comparison between TWAS, AS, and TWAS+AS microbiomes. TWAS_AS produces the highest amount of methane and most of this methane is coming from methane production for CO2 and H2 according to the 16s analysis pipeline. On the other hand, AS has the lowest amount of methane producers, but higher concentration of acetate causes higher methane production compared to TWAS and this matches the experimental data provided in [67].
